# Limitation of amino acid availability by bacterial populations during enhanced colitis in IBD mouse model

**DOI:** 10.1101/2022.10.03.510649

**Authors:** Tanner G. Richie, Leah Heeren, Abigail Kamke, Sophia Pogranichniy, Kourtney Monk, Trey Summers, Hallie Wiechman, Qinghong Ran, Soumyadev Sarkar, Brandon L. Plattner, Sonny T. M. Lee

## Abstract

Members of the Enterobacteriaceae family including *Escherichia coli* are associated with persistent gut inflammation during disorders like inflammatory bowel disease. This is due to rapid microbial colonization during dysbiosis combined with pathogenic tendencies. We characterized the dysbiotic gut community, defined potential functional pathways, and investigated crosstalk between host gene expression and microbial detections in an intestinal inflammation murine model. Members of Enterobacteriaceae family and the *Enterococcus* genus were highly detected in dysbiotic mice. These metagenome assembled genomes (MAGs) contained several virulence factors and metabolic pathways necessary to drive perpetual inflammation. Two Enterobacteriaceae MAGs with L-cysteine and L-taurine dioxygenases were strongly correlated with upregulation of the host gene CSAD, responsible for cysteine metabolism. Suggesting these bacteria compete with the host to utilize essential amino acids. We observed that bacterial isolates from dysbiotic mice displayed increased growth rates supplemented with L-cysteine, confirming that these microbes can utilize host nutrients to sustain inflammation.

**In Brief:** Inflammatory bowel disease is associated with an increase in Enterobacteriaceae and *Enterococcus* species, however the mechanisms are unclear. Richie *et al*. show that these bacterial populations use sulfur metabolism and tolerate host-derived immune-response, to drive host inflammation and fuel growth in the dysbiotic colon. Cultured isolates from dysbiotic mice indicated faster growth supplemented with L-cysteine, showing these microbes can utilize these essential host nutrients.

**Highlights:**

- Mice receiving native microbial FMT showed lower colon inflammation scores, higher microbial diversity, detections and gene expression similar to control mice.
- Dysbiotic mice displayed increased colon inflammation, higher detection of potential pathogenic MAGs, and upregulation of cysteine dioxygenase and other inflammation response genes
- MAGs assigned to *Enterococcus* and Enterobacteriaceae species were more frequently detected in dysbiotic mice, while almost absent in mice receiving FMT or control mice, they also contain several virulence factors and antibiotic resistance genes.
- These MAGs also display potential functions of utilizing host products and nutrients including nitrate, cysteine, and taurine to further fuel their growth and metabolism, which results in persistent host intestinal inflammation.
- Isolates in the Enterobacteriaceae family from dysbiotic mice utilize L-cysteine for growth, whereas isolates from FMT and control mice show no significant difference, indicating these bacteria can utilize the host derived cysteine.

## Introduction

Inflammatory bowel diseases (IBDs) are a category of chronic immune-mediated disorders with a wide range of outcomes and an unresolved etiology. The incidence of IBDs are increasing globally, with over 6 million cases, significantly increased number of diagnoses reported within the last 3 decades (1). The tri-factor of genetic, environmental, and microbial components are suspected to each play a significant role in the development of these complex disorders (2–4). A microbial cause for IBD has been suspected as far back as the mid-19th century when the disease was first described by Samuel Wilks in 1859 (5). While gut dysbiosis is associated with IBD, it has been difficult to definitively determine if these changes observed are causal or merely a consequence of IBDs. It is possible that dysbiosis initiates or contributes to the progression of IBD by amplifying and sustaining the local or systemic inflammatory process.

There is no consensus that IBDs are caused by classical pathogens. Recent studies show that commensal microbes can display pathogenic properties given the right circumstances, context and opportunity. These microbes have been called “pathobionts” (6–8). Specific microbes have been hypothesized to play a role in the development or aggravation of IBDs (9–11). Adherent invasive *Escherichia coli* (AIEC) and *Enterococcus faecalis* have been observed in IBD and Crohn’s disease (CD) patients that display virulence factors at higher rates compared to healthy patients (10–12). However, we still have limited understanding of the interactions between these bacteria and the host during the onset of IBD and related gastrointestinal inflammatory disorders. In addition to identification of relevant potential causal microbes of IBD, function of these microbes before and during disease, and how they interact with the host are crucial in understanding this complex disease dynamic. For example, *Akkermansia muciniphila* and butyrate-producing *Clostridia* have been shown to improve gut physiology and even alleviate symptoms of IBD and inflammation in both mice and humans (13–15). On the other hand, AIEC has been proposed as a possible cause of IBD, and studies have also found an increased number of virulent *E. coli* strains isolated from IBD patients (10, 16). Some studies suggest that the accumulation of AIEC and other *E. coli* strains in the gut is a consequence of inflammation, while other studies show that AIEC infection induces changes in the gut microbiota (9, 17). It is speculated that certain metabolic pathways of AIEC, such as utilization of host derived nitrate and uptake of sulfur rich amino acids like cysteine and taurine (7, 18–20), drive chronic inflammation within the gut environment. Currently, the mechanisms of inflammation associated with bacterial populations in the dysbiotic remain unclear.

In IBD patients, the chronically inflamed small intestine and colon are subjected to considerable oxidative stress (21, 22). Amino acid metabolism plays an important role in the nutrition and physiology of the host, and may influence the progress of IBD. Studies demonstrated that amino acids are critical for mucosal wound healing and are used as energy substrates of enterocytes. These amino acids may act to reduce inflammation, oxidative stress, and proinflammatory cytokines. Although experimental studies investigating the therapeutic efficacy of amino acids are promising, some clinical studies with supplementation in IBD patients are disappointing, lacking high-resolution in both host and microbial responses. Conflicting results warrant additional research with a goal of a deeper understanding how amino acid metabolism affects the underlying mechanisms of IBD.

In this study, we sequenced and assembled high-quality genomes of bacterial populations for system-wide simultaneous analysis of host gene expression and microbial functions. To focus on novel host-pathogen interactions that might reveal microbial impacts on the pathogenesis of IBD, we analyzed microbial functions and metabolic pathways that were complemented by the host. Using a combination of bioinformatic, genomic, and biochemical approaches, we highlighted biosynthesis and degradation pathways of cysteine and taurine in Enterobacteriaceae and *Enterococcus* populations that might contribute to initiation, amplification and sustaining intestinal inflammation in dysbiotic IL-10 KO mice. Protection from inflammation and colon damage was demonstrated in control and FMT groups with lower histologic inflammation scores and several immunologic indicators. Importantly, we also showed that Enterobacteriaceae populations isolated from mice displayed growth specifically by utilization of L-cysteine. Our experiments further our fundamental understanding of the impact of host-microbe interaction on intestinal inflammation, showed that the understanding of the congruent expressions of host gene expressions and microbial functions represent a robust way to discover potential implications of pathobionts, and thereby provides a target for the generation of novel microbial treatment options for IBD.

## Results & Discussion

Fecal content from pups were collected at the final week (week 24) of the study. Genomic DNA was extracted, and shotgun sequencing was conducted (Figure 1). Briefly, mouse pups displayed vertically transmitted dysbiosis from cefoperazone antibiotic exposure, except for the control group (control/no-CPZ), and received either FMT (CPZ/FMT), saline gavage (CPZ/gavage-with-PBS), or no gavage (CPZ/no-gavage) during and after development. We employed genome-resolved metagenomics for an in-depth characterization of the gut microbiota, and combined with host response markers - colon RNA sequencing, serum immune markers and histology to provide insights into the pathogenesis of the microbiota.

**Figure 1.**
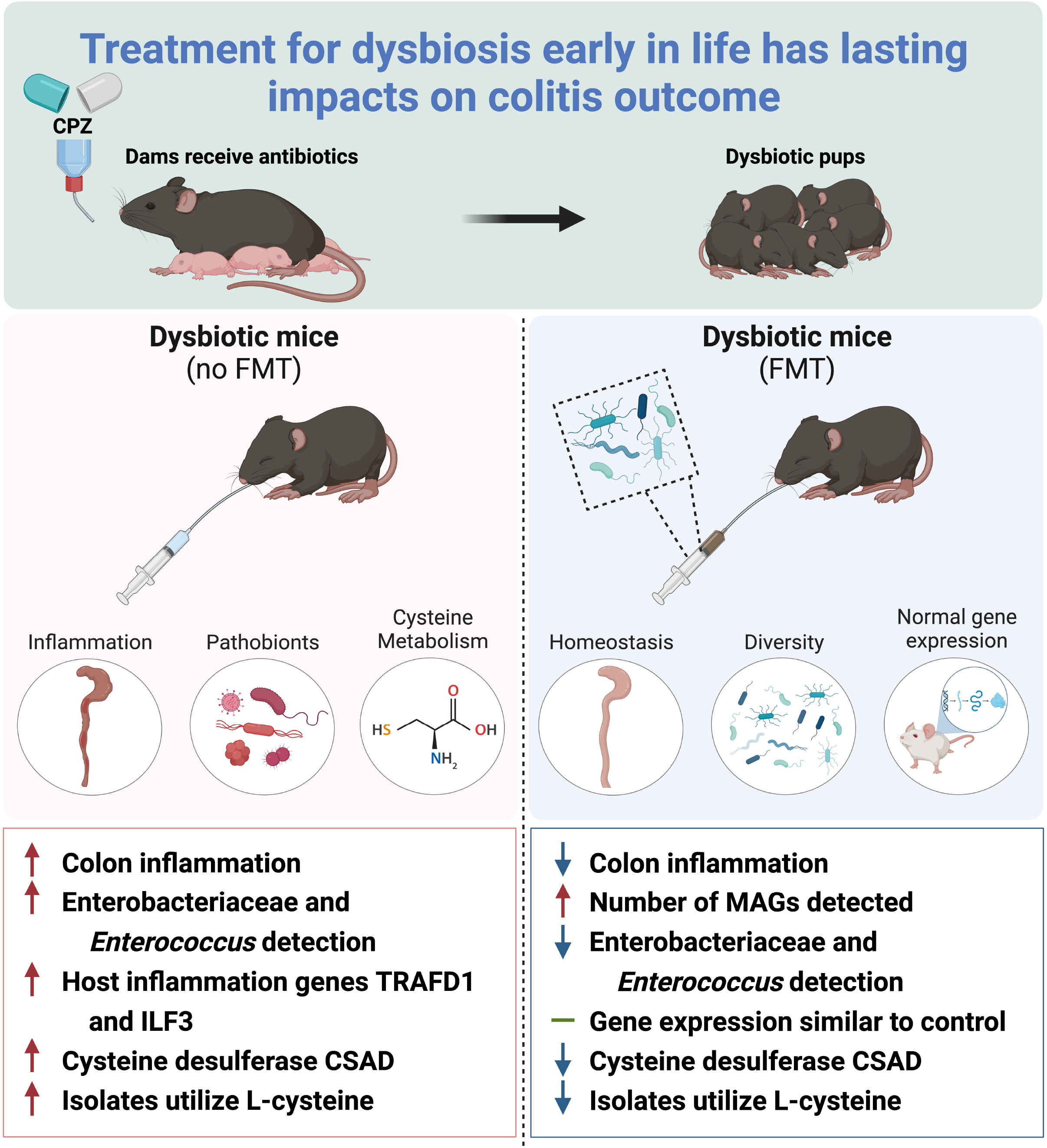
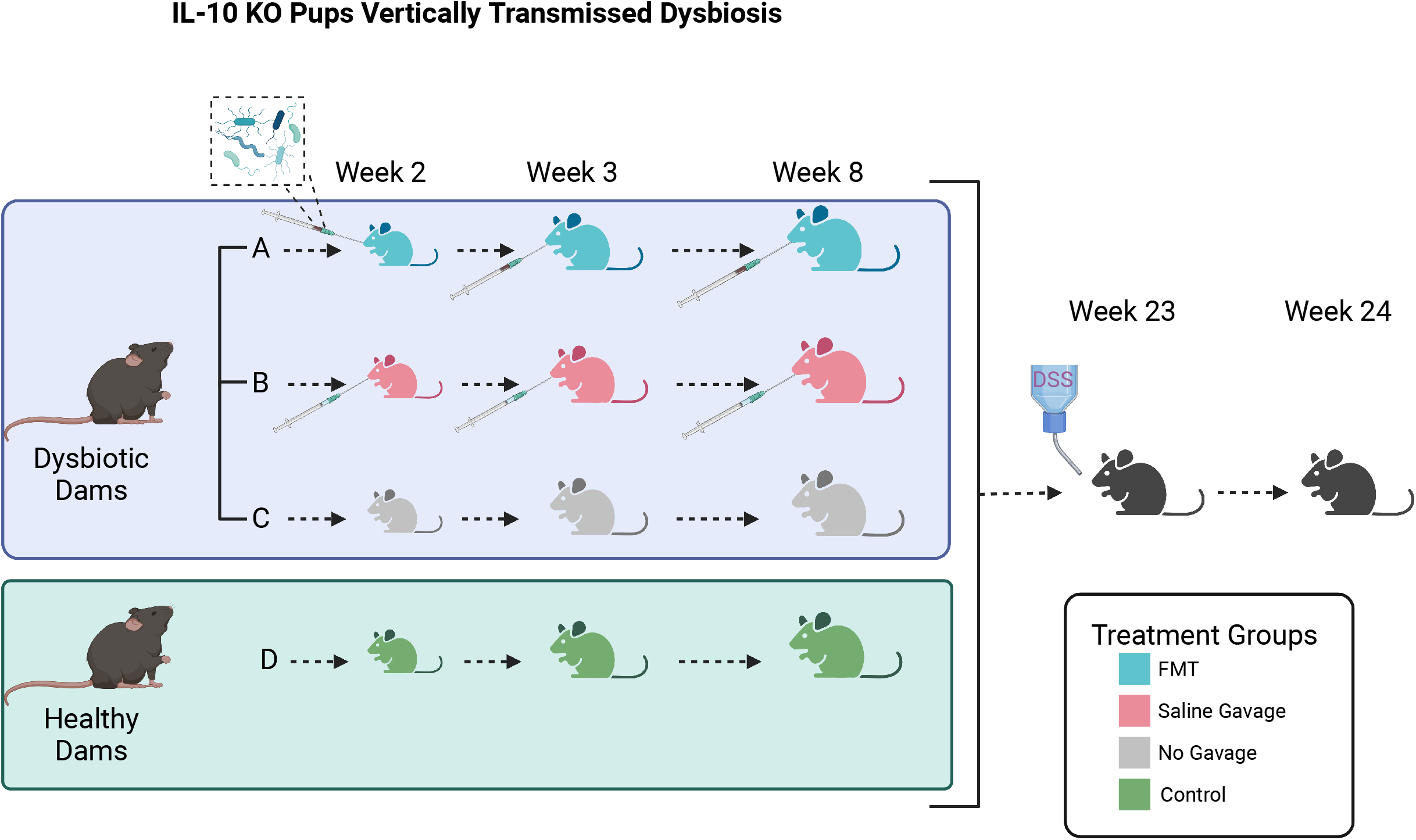
Graphic of experimental design highlighting crucial conditions and the timeline by inclusion of important dates of the experiment and defining the treatment groups.

Shotgun sequencing of the fecal samples resulted in a total of 623,630,018 sequences with an average of 12,228,040 ± 1,646,545 reads per metagenome (Supplementary Table S1). Coassemblies of the 4 experimental group (n= 51; control/no-CPZ, n= 19; CPZ/no-gavage, n= 9; CPZ/gavage-with-PBS, n= 13; CPZ/FMT, n= 10) metagenomes resulted in an average of 90,779 ± 14,373 contigs that were longer than 1,000 nucleotides.

### FMT resulted in reduced colon inflammation and changes in systemic serum markers

We investigated the host response to FMT to gain a perspective of the impacts later in life from reintroducing native microbes during development of the neonates. We noticed that mice that received FMT displayed less colon surface damage, fewer lamina propria infiltrating lymphocytes, and significantly lower inflammation scores than dysbiotic mice that did not receive FMT (p<0.02). Mice that received saline gavage or no gavage displayed the highest histopathologic score with evidence of epithelial damage, lymphocyte infiltration, and loss of goblet cells (p<0.029) (Figure 2A, Supplementary Table S2). Select systemic serum markers indicative of broad-scale inflammation were measured in serum of IL-10 KO mice. The cytokine panel showed low level trends for the control mice compared to other groups (Figure 2B, Supplementary Table S2). Notably, FMT mice resulted in significantly higher levels G-CSF (p=0.0011), IL-12p70 (p=0.044), IL-17 (p=0.025), TNFα (p=0.0403), and IL-6 (p=0.0028) compared with control mice, suggesting that certain beneficial microbes, when acquired early in life, play a deterministic role in host immune modulation and priming, as well as maintaining homeostasis (23–25). One cytokine, IL-2, displayed lower levels in dysbiotic mice compared to FMT-treated mice; however, IL-2 was not significantly different from control mice (p<0.049, Supplementary Table S2). With few significant increases in common IBD inflammation indicators like IL-1β and IL-17 (26, 27), we observed little evidence of a systemic, uncontrolled, immune response, regardless of treatment group. In summary, IL-10 KO mice receiving FMT displayed less colon inflammation indicating a successful FMT with long term inflammation relief and cytokine modulation.

**Figure 2.**
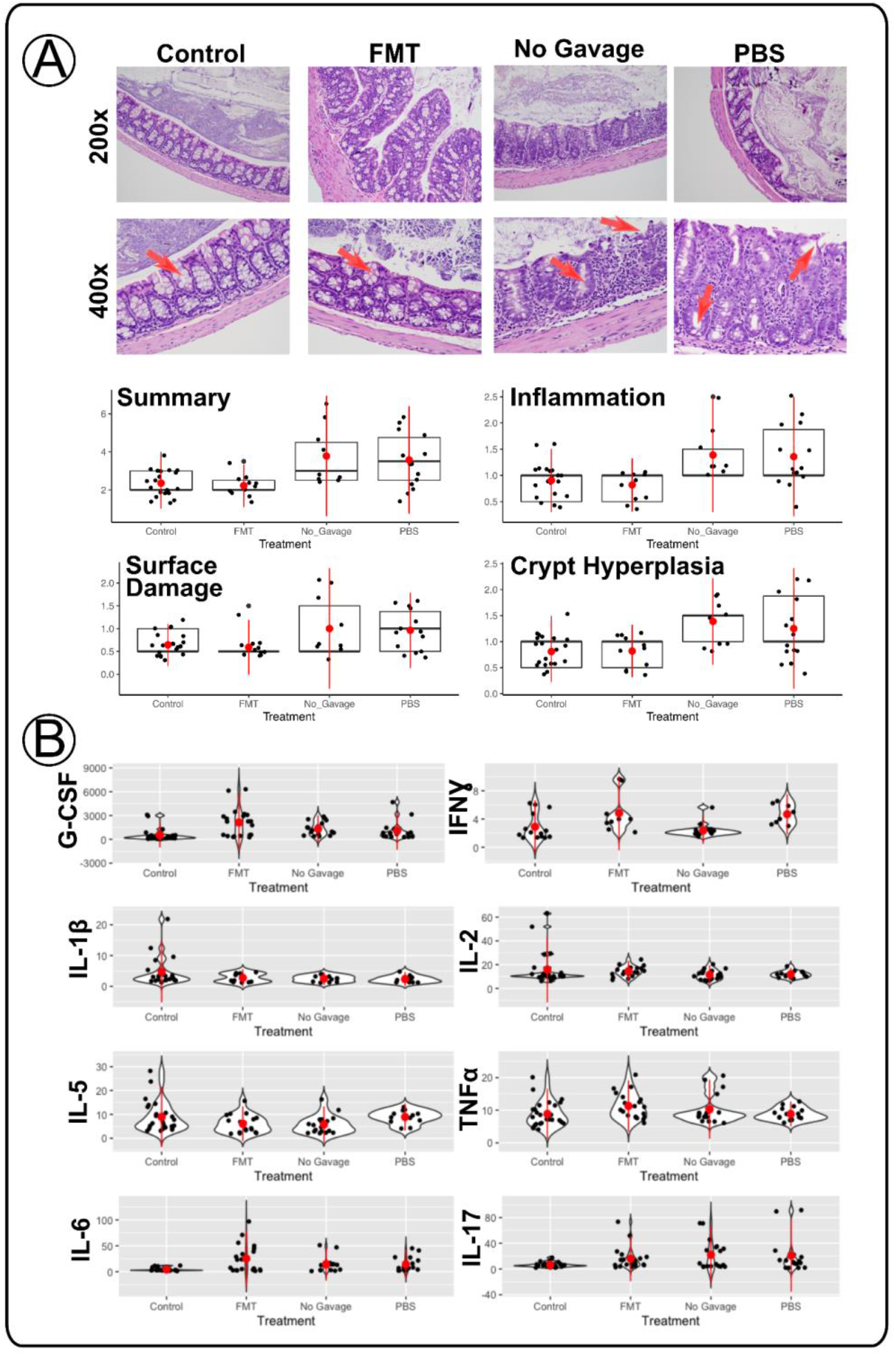
Dysbiosis drives persistent host response in colon inflammation with absence of large systemic response. (A) Histologic images of IL-10 KO colon representing each treatment group. Flags indicate sites of goblet cell status or surface damage present as indicated. Summary scores are shown for each treatment group including individual scores for inflammation, surface damage and crypt hyperplasia. (B) Pro-inflammatory cytokine measurements of IL-10 KO serum for cytokines relevant in IBD development.

### Upregulation of several IBD and cancer related genes in dysbiotic mice highlight importance of microbial biodiversity

To further investigate host response to long term dysbiosis, we performed RNA sequencing on mice colon samples (n=3) from each treatment group. First, we performed a Gene Ontology (GO) Analysis to gain insight into broad gene categories that were disproportionately upregulated or downregulated by treatment group comparisons. Six pairwise comparisons revealed gene regulation changes between treatment groups (Supplementary Figure S1-S6). Comparisons between control and FMT mice showed only 6 gene categories significantly different, (3 upregulated-, 3 downregulated-classes), while significantly more differences were seen with comparisons between FMT and the CPZ/gavage-with-PBS (35 upregulated, 32 downregulated; 67 total), as well as FMT compared with CPZ/no-gavage (84 upregulated, 45 downregulated; 129 total). Similarly, comparisons between control mice and CPZ/gavage-with-PBS (20 upregulated, 34 downregulated; 54 total), and the CPZ/no-gavage groups (59 upregulated, 29 down; 88 total) also showed several categorical differences. We observed significant differences in several gene families between dysbiotic and FMT mice groups, including multiple inflammatory cytokine/chemokine activity, sulfur dioxygenase, sulfur transport activity, cysteine receptor binding, and nitric oxide synthase binding. Nearly identical trends in gene family differences between control mice and dysbiotic groups were observed. Finally, differences between FMT and control groups were fewer, with significant differences noted in gap junction channel activity, cytokine/chemokine activity, IL-17 receptor activity, and sulfur compound binding, all suggesting the FMT successfully reduced gut inflammation in the host. We examined these gene families because of their known involvement in the development or more severe symptoms of IBD in human patients (20, 26, 28, 29). As discussed previously, cytokine and chemokine modulators initiate inflammation during an immune response, with IFNƔ, IL-6, and IL-17 acting as early indicators of IBD (27). Sulfur dioxygenase activity is a precursor step in the production of cysteine, along with the transporter genes to gain access to more sulfur ions to construct cysteine (30). In IBD patients, low cysteine and taurine levels have been observed, and pathogenic enteric bacteria such as *Escherichia coli* likely benefit from sulfur metabolism (19, 20), indicating from our data that dysbiotic mice might have lower cysteine levels contributing to the upregulation of cysteine biosynthesis gene families. Nitric oxide is also a crucial component of the host immune response to potential pathogens. Nitric oxide is generally toxic to gut microbes (31), is known to be significantly increased in dysbiotic mice, which suggests that its presence indicates a pro-inflammatory immune response in the gut, and is thus another potential biomarker of active IBD (32).

To gain more insights into the host expressed gene functions, we examined fold changes of individual genes in treatment group comparisons (Figure 3, Supplementary Table S3). We observed several upregulated genes associated with colitis and IBD development, including *ABCA2*, *Acss1*, *Araf*, *CSAD*, *Ilf3*, and *TRAFD1*, showing similar fold change patterns across treatment groups. Each of these genes were upregulated in dysbiotic mice as compared to the CPZ/FMT and control/no-CPZ groups. These genes function as an ABC transporter (*ABCA2*) (33, 34), a member of the tricarboxylic acid cycle (*Acss1*), growth promoter kinase (*Araf*) (35), cysteine desulfurase (*CSAD*), interleukin binding enhancer (*Ilf3*) (36), and a negative regulator of the innate immune response against pathogens (*TRAFD1*) (37, 38) that are found to be upregulated in IBD and colon cancer development. *TRAFD1* upregulation suggests a host attempt to slow immune response common during pathogen infection response. *CSAD* gene upregulation seen in the dysbiotic groups indicated that the host was increasingly utilizing cysteine as a sulfur source; a common finding in IBD patients is low serum cysteine levels (39), which could indicated the dysbiotic mice in our study were experiencing low cysteine levels as well. Formation of homocysteine occurs during biosynthesis of cysteine, and an increase in homocysteine levels has been shown in IBD patients, indicating that the host might be starved of cysteine (40, 41). The genes accounted for here were also represented above in the gene family categories that were differentially regulated in dysbiotic mice compared to FMT and control mice. Our results collectively suggest that these genes are an indication of a hosts’ response to dysbiosis in the gut, and its immune responses to regulate inflammation

**Figure 3.**
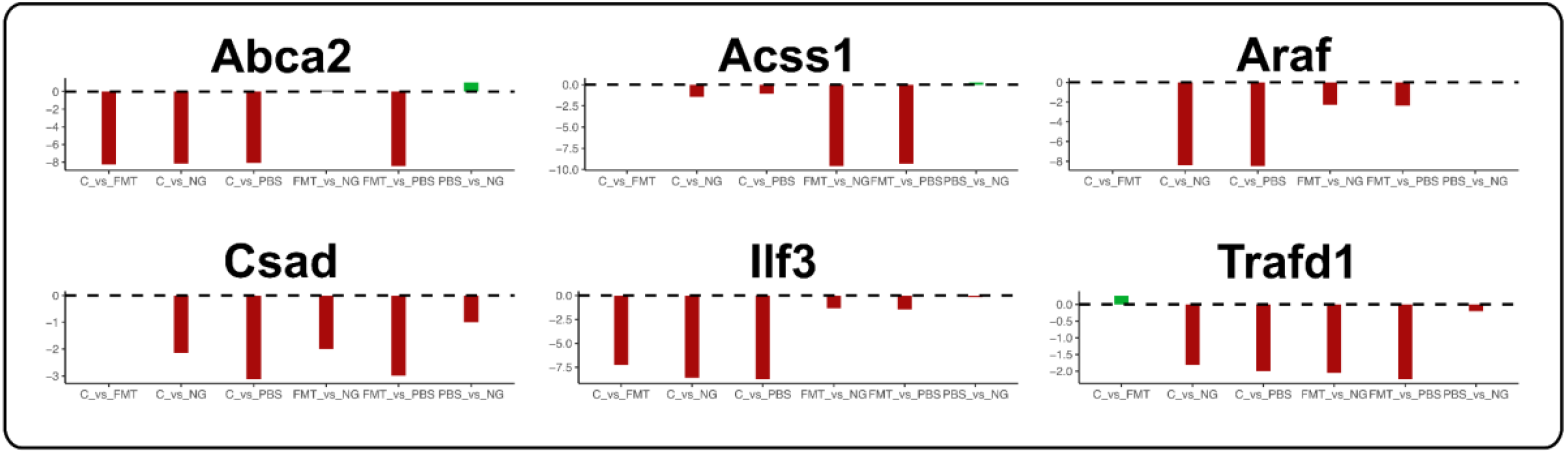
IBD and cancer associated genes upregulated in dysbiotic IL-10 KO mice. Fold change plots by treatment group comparisons.

### Distinct bacterial populations are associated with inherently dysbiotic IL-10 KO pups

Due to changes in certain bacterial populations, conditions in dysbiosis-associated gut are reported to result in chronic persistent intestinal inflammation (16, 42, 43). Therefore, with the knowledge of the hosts’ gene expressions, we next examined the effect of reintroducing native intestinal microbial communities via FMT early in life on gut microbial composition postdevelopment. We reconstructed metagenome-assembled genomes (MAGs) that were >70% complete with <10% redundancy as predicted by bacterial single-copy core genes (44, 45), from the co-assembled treatment group metagenomes (46, 47). We resolved a total of 677 MAGs, and then removed redundancy by selecting a single representative for each set of genomes, resulting in 190 non-redundant MAGs (Supplementary Table S4) from the 4 experimental groups. There was a large representation of Firmicutes (n=178) with Proteobacteria (n=6), Bacteroidota (n=1), Actinobacteria (n=3) and Verrucomicrobiota (n=2) having lower detection. At the class level, Clostridia (n=158) was the most dominant, followed by Bacilli (n=17), Gammaproteobacteria (n=3), Coriobacteriia (n=3), Negativicutes (n=1), Dehalobacteriia (n=1), Bacteroidia (n=1), Verrucomicrobia (n=1) and five remained unidentified (Figure 4A).

**Figure 4.**
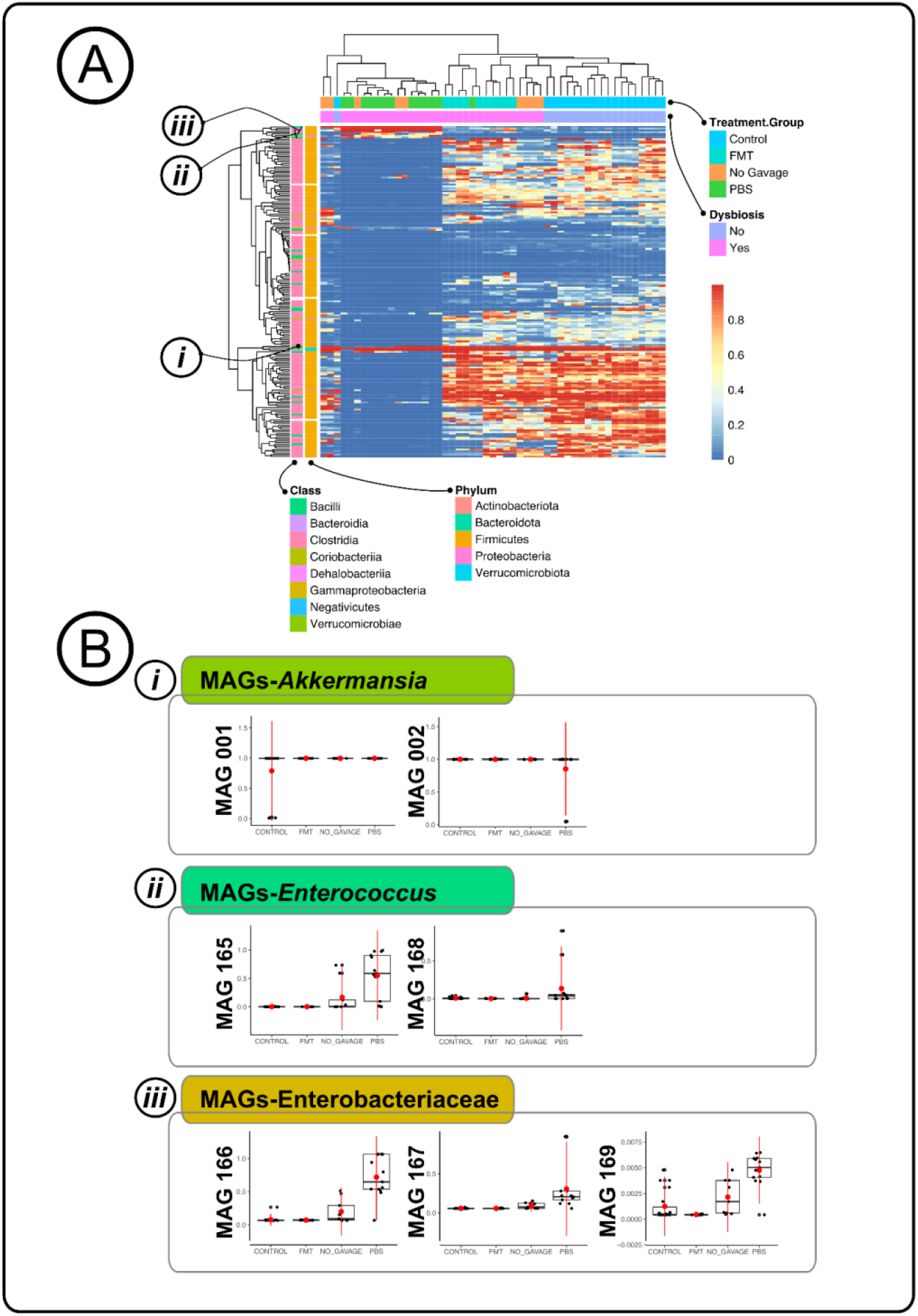
Distinct bacterial populations that were associated with inherently dysbiotic IL-10 KO pups. (A) Clustered heatmap of IL-10 KO with the non-redundant MAGs. (B) Boxplots of detection values of specific MAGs of interest, *Akkermansia, Enterococcus*, and Enterobacteriaceae by treatment group.

We observed that the gut microbiota of mice that received FMT resembled that of mice that were not dysbiotic. Dysbiotic mice that received either gavage of PBS or no gavage were distinctly different from non-dysbiotic and FMT mice (Figure 4A, Supplementary Table S5). We used the MAG’s detection values to calculate the Bray-Curtis dissimilarity and showed that there was a significant clustering of control/no-CPZ and CPZ/FMT group (Figure 4A). We also observed a wider clustering pattern in the CPZ/no-gavage and CPZ/gavage-with-PBS groups. The differences among the treatment groups were mainly attributed to the absence and presence of the families Enterococcaceae and Enterobacteriaceae which exhibited the largest differences in MAG’s detection (Supplementary Table S5). Our results corroborated with other studies that showed an increase in relative abundance of bacterial populations from families Enterococcaceae and Enterobacteriaceae (10, 16, 48, 49).

Although previous studies have identified changes in relative abundance of Enterococcaceae and Enterobacteriaceae bacterial populations, it has been challenging to identify the exact culprit causing gut inflammation due to resolution of the studies (50, 51). In our study, of the 190 non-redundant MAGs, we resolved to a high-resolution level, 7 MAGs that were of interest-MAGs 001 and 002 (hereafter referred as MAGs-*Akkermansia*), MAGs 165 and 168 (hereafter referred as *MAGs-Enterococcus*), and MAGs 166, 167, and 169 (hereafter referred as MAGs-Enterobacteriaceae). We resolved MAGs-*Akkermansia* to their respective species level (MAG 001: *Akkermansia muciniphila* and MAG 002: *CAG-485 sp002362485*), while MAGs-*Enterococcus* were also annotated to the species level (MAG 165: *Enterococcus faecium* and MAG168: *Enterococcus faecalis*). While assigning identity to MAG 166 (Enterobacteriaceae), and MAG 167 (Enterobacteriaceae) was challenging due to the highly heterogeneity among the bacterial populations in this family, we managed to resolve both MAGs to a 100% completion (MAG 166 - redundancy value 5.6%, and MAG 167 - redundancy value 1.4%), obtaining valuable MAG genomic information for downstream analyses. In addition, MAG 169 was assigned to the species *Proteus mirabilis* (completion 100%, redundancy 0%). Detections of both *Enterococcus* and Enterobacteriaceae MAGs by treatment group indicated a lower presence in control/no-CPZ and mice that receive FMT, with significantly higher detections in the dysbiotic mice with either the PBS gavage or no gavage (Figure 4B). We surmised that MAGs-*Enterococcus* and MAGs-Enterobacteriaceae possessed functions that not only enable them to proliferate and outcompete other microbial competitors during dysbiosis, but may also have contributed to persistent gut inflammation in the host.

We used Pathosystems Resource Integration Center (PATRIC) web portal (52, 53) and constructed phylogenetic trees to further verify the identity of the MAGs. We verified that MAG 001 annotated to *Akkermansia muciniphila*, and MAG 002 was closely related to the *Barnesiella* genus in the Bacteriodota phylum. The other 5 MAGs were from two different taxonomic groups - *Enterococcus* (MAG 165 and 168) with the closest relationship to *Enterococcus faecium* and *Enterococcus faecalis* respectively, and Enterobacteriacaea (MAGs 166, 167, and 169) had the closest phylogenetic relationship with *Enterobacter, Kosakonia*, and *Proteus*, respectively (Supplementary Figure S7-S13). Out of the three groups of MAGs of interest, *MAGs-Akkermansia* was highly detected in all of the samples (control/no-CPZ: 0.894 ± 0.0496; CPZ/no-gavage: 0.997 ± 0.0005; CPZ/gavage-with-PBS: 0.924 ± 0.0505; CPZ/FMT: 0.998 ± 0.0005), regardless of whether the mice were dysbiotic or not. Although control mice had significantly lower MAG 001 detection compared with other groups (p<0.0421, Figure 4B), higher detection of MAG 001 was observed in all treatment groups. MAGs-*Enterococcus* and MAGs-Enterobacteriaceae had a very different detection pattern as that of *MAGs-Akkermansia* (Figure 4B). Both MAGs-*Enterococcus* and MAGs-Enterobacteriaceae displayed low detection in the control/no-CPZ (MAGs-*Enterococcus:* 0.005 ± 0.0015, MAGs-Enterobacteriaceae: 0.005 ± 0.0035) and CPZ/FMT (MAGs-*Enterococcus:* 0.002 ± 0.0005, MAGs-Enterobacteriaceae: 0.003 ± 0.0006) groups, however, were highly detected in the CPZ/no-gavage (*MAGs-Enterococcus:* 0.087 ± 0.0501, MAGs-Enterobacteriaceae: 0.057 ± 0.0226) and CPZ/gavage-with-PBS (MAGs-*Enterococcus:* 0.343 ± 0.0776, MAGs-Enterobacteriaceae: 0.303 ± 0.0583) group. We also observed a high statistical significant difference in the detection of MAGs-*Enterococcus* and MAGs-Enterobacteriaceae between mice that were treated with FMT as compared to mice received PBS or were not given any gavages: MAG 165 (FMT/PBS t-test, t statistic= −4.99, p=0.0003, FMT/no-gavage t-test, t statistic= −1.71, p=0.12, PBS/control t-test, t statistic= 4.98, p=0.0003, FMT/control t-test, t statistic= −0.93, p=0.36, no gavage/control t-test, t statistic= 1.7, p=0.13, Figure 4B); MAGs 166 and 167 had a similar pattern, with FMT and control groups having significantly lower detections than dysbiotic groups (p<0.027, no significance between FMT and control). Finally, MAG 169 was significantly lower in FMT compared to all groups, including control mice (p<0.027 Figure 4B).

### Coherence between host and microbe highlighted attributes to worsened outcomes in IBD

Identifying host-microbe interactions during dysbiosis-associated intestinal inflammation can provide insight into both the host and microbial pathways that affect inflammation outcomes. So far, we identified specific host modulations in response to FMT, as well as specific bacterial populations that were associated with the dysbiosis. To further determine likely interactions between host and microbe modulations, we employed a discriminant analysis using the mixOmics package encoding DIABLO in R (54). We used a correlation cutoff at 0.85 to ensure robust and biologically relevant linkages between host genes and MAGs (Figure 5A). We demonstrated that Enterobacteriaceae MAGs were positively correlated with inflammatory and host amino acid metabolic response genes, while the Lachnospiraceae and Oscillospiraceae families were negatively correlated with inflammatory genes. Our analysis suggested that MAGs from the Enterobacteriaceae family might drive host response in a dysbiotic gut through crosstalk with the host immune system, and uptake of essential amino acids. We showed that two host genes displayed both positive and negative correlations between a set of MAGs; *CSAD* and *TRAFD1*. *CSAD*, as shown above to be upregulated in dysbiotic mice, was positively correlated to two MAGs: Enterobacteriaceae MAGs 167 and 169. This gene was also negatively correlated with several MAGs closely matching the families Lachnospiraceae and Oscillospiraceae. The gene *CSAD* catalyzes the decarboxylation of cysteine sulfinate to hypotaurine, eventually to taurine (55), with cysteine shown to be essential in the recovery of the host from chronic colon inflammation and reduced IBD symptoms (56, 57). Thus, the strong positive correlation between *CSAD* and MAGs-Enterobacteriaceae suggests that these MAGs may be competing with the host for sulfur-rich amino acids cysteine and taurine as an alternate metabolic pathway to fuel growth leading to over colonization of these pathobionts in the gut (19, 20). The *TRAFD1* gene, responsible for controlling immune response to pathogens, was positively correlated with Enterobacteriaceae MAG 169, and negatively correlated with multiple MAGs from the Lachnospiraceae and Oscillospiraceae families. One MAG of the *Dorea* genus displayed several positive correlations to the *Fut10*, *Wrap53*, *Primpol*, *Chd5*, and *Gnl1* genes. Collectively we were able to identify coherent patterns between the host gene expression and microbial populations, and showed potential metabolic pathways these MAGs could utilize to drive host response, while fueling growth and metabolism of these pathobionts.

**Figure 5.**
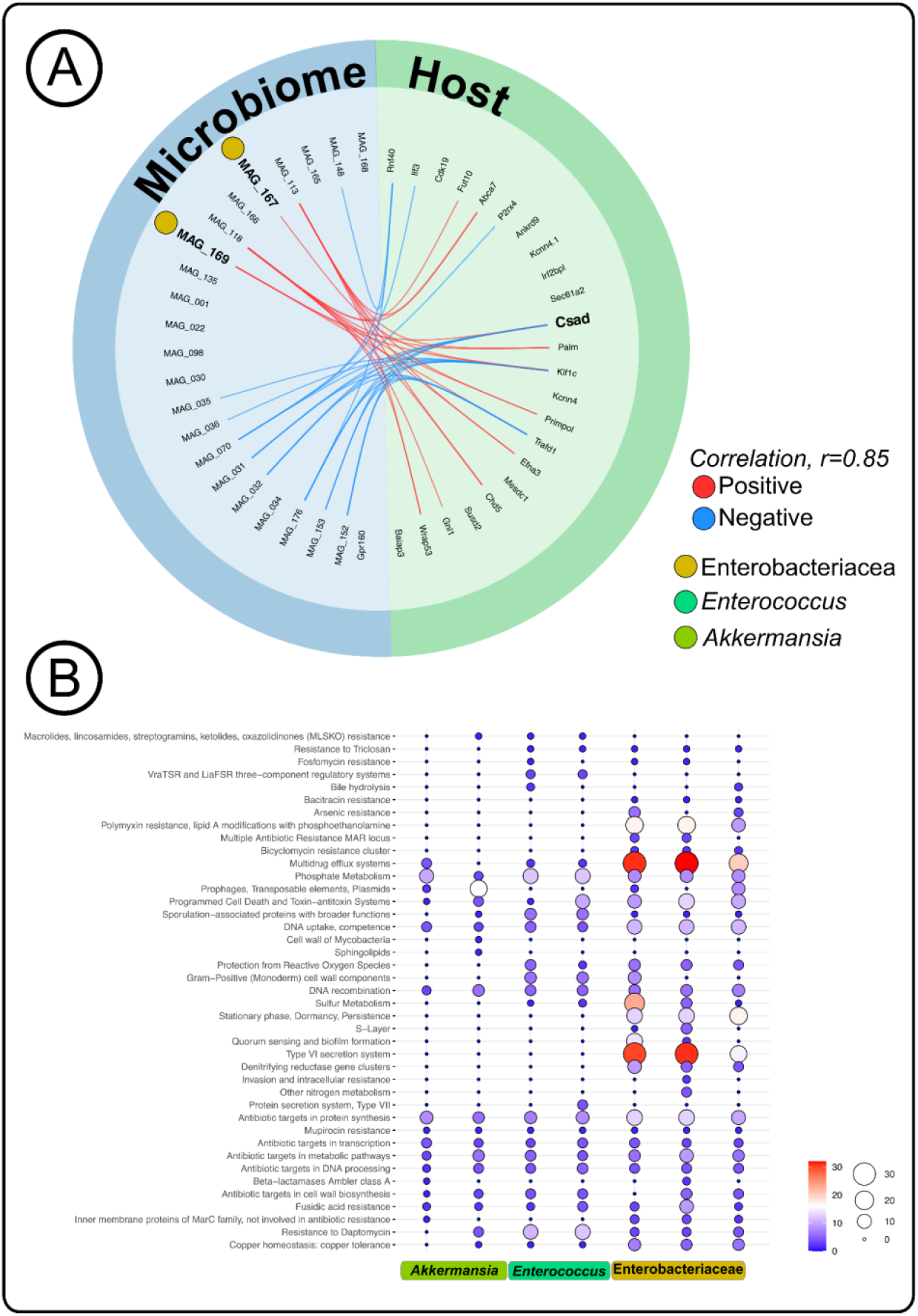
Coherence between host and microbe highlighted attributes to worsened outcomes in IBD. (A) Correlation analysis of non-redundant MAG detection values with RNA sequencing counts for a subset of host genes. (B) Pathway analysis of virulence gene counts for each MAG of interest, *Akkermansia, Enterococcus*, and Enterobacteriaceae.

### Utilization capability of amino acids by bacterial members impacted host IBD outcomes

We used discriminant analysis and showed that MAGs-*Enterococcus* and MAGs-Enterobacteriaceae detection correlated with a negative impact on host gene expression, resulting in persistent intestinal inflammation. Several previous studies have indicated that the development of IBD could be associated with microbial factors including pathways driving host inflammation in addition to sustaining colonization, virulence factors such as secretion systems and effectors, and antibiotic resistance genes (9–11). Therefore, we assessed potential gene functions in the 7 MAGs of interest (MAGs-*Akkermansia*, MAGs-*Enterococcus*, and MAGs-Enterobacteriaceae) to examine whether genome level alterations related to those functions were observed in our resolved MAGs (Figure 5B). We observed an absence of virulence genes that were detected in *MAGs-Akkermansia;* however, multiple antimicrobial resistance genes were also absent. Previously, *Akkermansia muciniphila* has been implicated as a commensal member of the gut microbiota, as a mucus degrader, and has been associated with reduced overall colon inflammation, which may be due to its role in strengthening tight junctions of intestinal epithelial cells (14, 58, 59). MAGs-Enterobacteriaceae clustered on its own branch with highly detected denitrifying gene clusters, T6SS, and multidrug efflux pumps (Figure 6A). Our results along with other studies, found a relative increase in virulent Enterobacteriaceae bacterial populations in IBD patients (16, 60). Our results indicated that the high detection of MAGs-Enterobacteriaceae in dysbiotic mice resulted in more severe intestinal inflammation in these mice, and MAGs-Enterobacteriaceae possessed the potential functions to exacerbate the inflamed gut. In addition, the MAGs-Enterobacteriaceae group included a large presence of nitrate and nitrite reductases capable of denitrifying nitrate and nitrite to ammonia (Figure 6A). COGG genes from both MAGs-Enterobacteriaceae and *MAGs-Enterococcus* groups included the *NarL* and *NarX* gene families, a two component system regulating various nitrate and nitrite reductases. The presence of nitrate and nitrite reductases demonstrated MAGs-Enterobacteriaceae’s ability to completely denitrify nitrate to ammonia, which is critical for quick growth under anaerobic conditions like the gut environment. Some of the MAGs in both groups *MAGs-Enterococcus* and MAGs-Enterobacteriaceae possessed sulfur metabolism functional potential (Figure 6B). The amino acids cysteine and taurine are utilized and broken down during both host and microbial sulfur metabolisms (61, 62). Previous studies have shown the importance of cysteine and taurine in host remediation of intestinal inflammation by providing regulation to pro-inflammatory cytokines, and relieving oxidative stress in addition to lumen endothelial cell dysfunction (56, 63).

**Figure 6.**
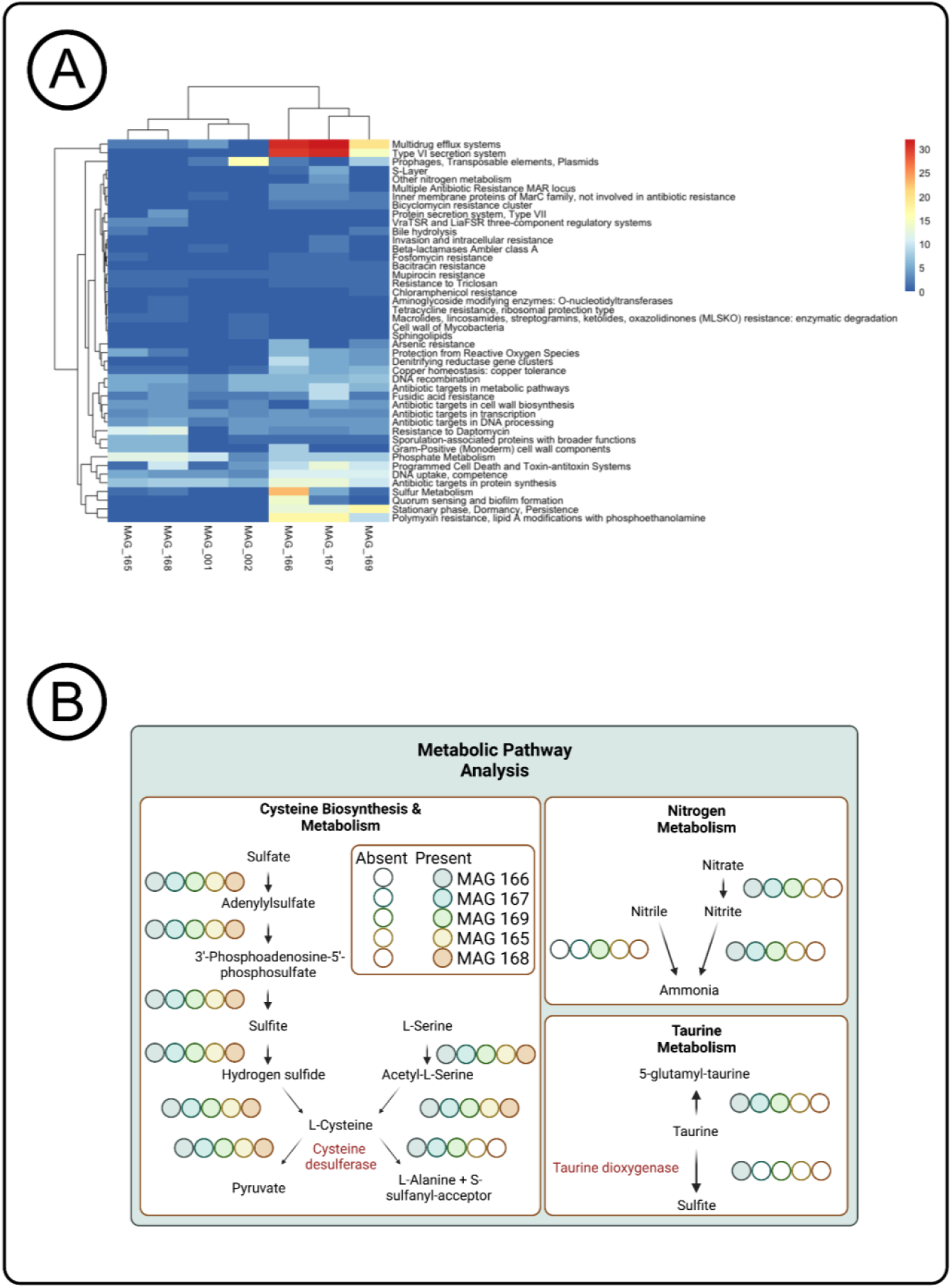
Utilization capability of amino acids by bacterial members impacted host IBD outcomes. (A) Clustering based on subsystem metabolic pathways of each MAG of interest *Enterococcus* and Enterobacteriaceae. (B) Metabolic pathway maps of each MAG of interest *Enterococcus* and Enterobacteriaceae and the genes in sulfur, taurine and nitrogen metabolisms that are present.

An important component of the host inflammatory immune responses is the ability of the host to reduce and limit the inflammation. We thus hypothesized that *MAGs-Enterococcus*, and MAGs-Enterobacteriaceae might limit the availability of cysteine and taurine to the host, contributing mechanistically to persistent intestinal inflammation. We analyzed potential metabolic pathways of MAGs-Enterobacteriaceae and *MAGs-Enterococcus* to understand the metabolic activity in these MAGs that coincided and competed with host metabolism. We used the KEGG pathway analysis and showed that MAGs-Enterobacteriaceae possessed complete desulfurization gene pathways, involving both biosynthesis and degradation of the amino acids cysteine and taurine, whereas *MAGs-Enterococcus* possessed complete metabolic pathways for cysteine alone (Figure 6A). All of the Enterobacteriaceae MAGs harbored two genes; *SufS*, a cysteine desulfurase and *TauD*, a taurine dioxygenase, that are essential for degradation of these two sulfur-containing amino acids by cleaving the sulfur group (64–66). *SufE*, cysteine desulfurase gene, was identified in *MAGs-Enterococcus* which breaks down cysteine similarly to Enterobacteriaceae (67). To further verify the utilization of cysteine in MAGs-Enterobacteriaceae, we cultured and isolated 31 unique Enterobacteriaceae bacteria colonies from CPZ/FMT, control/no-CPZ, CPZ/gavage-with-PBS, and CPZ/no-gavage mice fecal samples. Altogether, 8 unique isolates were cultured from mice receiving FMT, 8 from the CPZ/gavage-with-PBS group, 8 from the CPZ/no-gavage group and 7 from control/no-CPZ mice. We conducted nutrient dependency assays using these 31 bacterial populations with or without cysteine in M9 broth (Supplementary Table S6). Isolates grown in cysteine supplemented media resulted in a significantly higher growth curve than isolates in minimal media (Figure 7). Isolates cultured from the control and FMT groups showed no significant differences at any time point during the assay (Figure 7). Eight isolates from the no-gavage and gavage-with-PBS groups displayed significantly increased growth rate compared to the unsupplemented M9 broth (Figure 7). These results suggest that Enterobacteriaceae in dysbiotic mice were using cysteine more efficiently for growth than Enterobacteriaceae from healthy mice or mice that received FMT. This result along with the knowledge that the host experiences low cysteine levels highlights an important potential mechanism that these bacteria could shift to sulfur metabolism to fuel growth and host inflammation, which has been speculated (20, 39, 40). Our observation of highly detected MAGs-Enterobacteriaceae and *MAGs-Enterococcus* in the dysbiotic mice with inflamed gut, the identification of cysteine and taurine metabolizing genes in these bacterial populations along with showing use of cysteine for increased growth, combined with the corresponding decarboxylation of cysteine gene expressions in the host; suggests that MAGs-Enterobacteriaceae and MAGs-*Enterococcus* limits the availability of cysteine to the host which are essential for restoring gut immune homeostasis by attenuating inflammatory responses (19, 20, 39). Putting all together, our results indicated the potential for our resolved MAGs to utilize essential host nutrients for metabolism in addition to consuming host immune response products like nitrate and nitric oxide for promotion of growth. These functional mechanisms alongside the virulence factors identified provide an advantage for these MAGs to over colonize in the dysbiotic gut, which may contribute to persistent chronic intestinal inflammation in the host, driving disease symptoms and progression.

**Figure 7.**
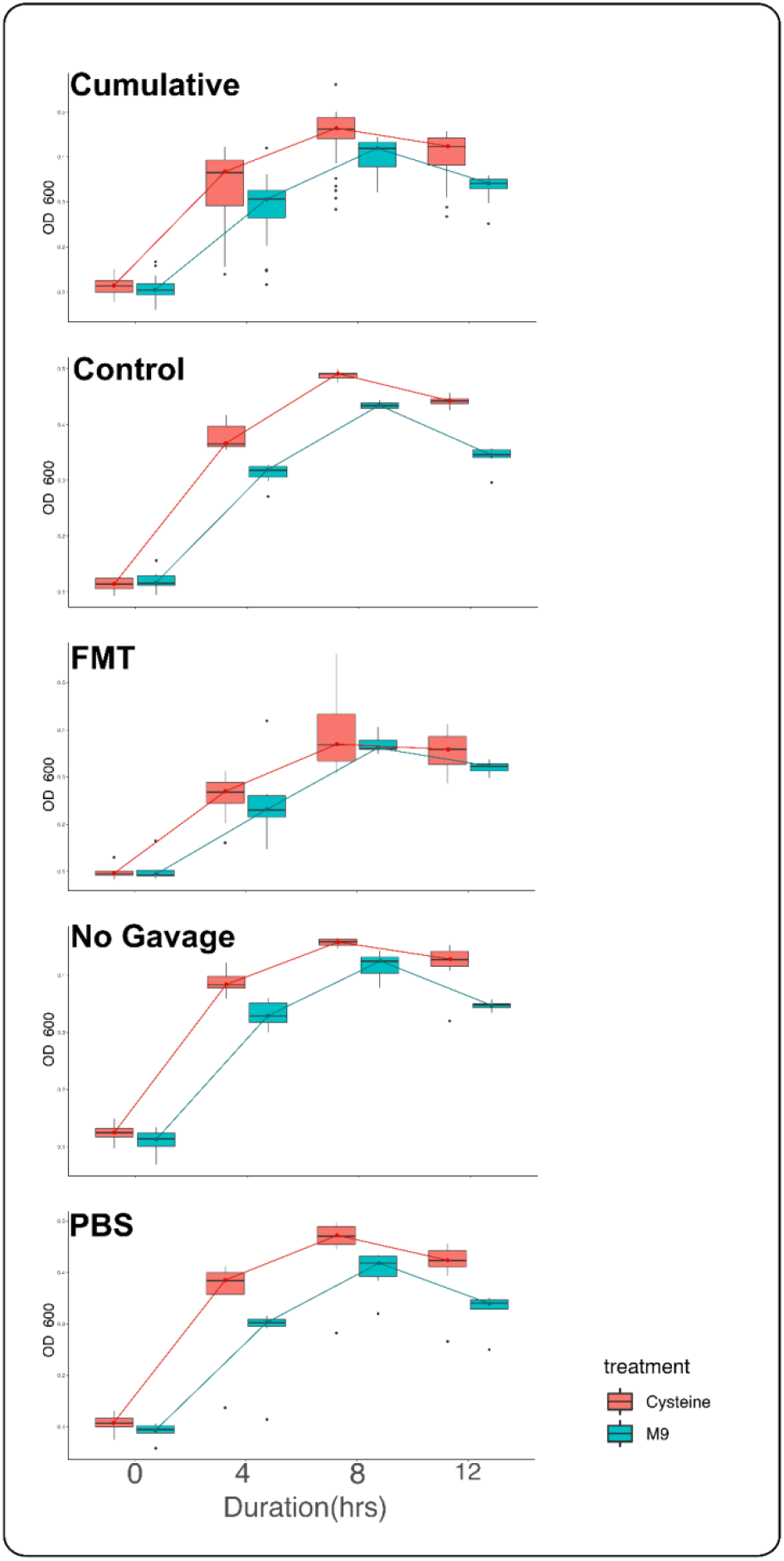
Growth curves of isolates cultured in cysteine and taurine from IL-10 KO mice in each treatment group.

Here, we use an array of host physiological and genetic data in addition to microbial metagenomic and metabolic information, for the first time, providing insights into possible mechanistic explanations of persistent intestinal inflammation in a dysbiotic gut. We showed a set of potential metabolic pathways that pathobionts are capable of utilizing to gain a foothold in the dysbiotic gut, driving persistent host inflammation, and leading to colitis. We demonstrated in our study that members of Enterobacteriaceae have the functional capabilities to compete with the host for L-cysteine as a sulfur source, while simultaneously driving inflammation with the products of this sulfur metabolism. Another well-known pathway involved with gut inflammation is nitrate metabolism (17, 68). The host produces nitrate and nitric oxide (26, 60) to ward off potential pathogens, however several Enterobacteriaceae members are tolerant and use one or both of these host response products (69). Essentially, the host inflammatory response fuels the growth of these pathobionts stimulating further immune modulation. These metabolic pathways could harbor mechanisms at which these bacteria use to transition from commensal to pathobiont (20). Thus switching to these alternate inflammation driving metabolisms, creating a chronic cycle of host inflammation and damage to the colon. While our findings provide a compelling link between the Enterobacteriaceae and *Enterococcus* MAGs and sulfur metabolism, these are potential functions and more work is needed to understand the true functional capacity in regards to sulfur metabolism. These potential functions were validated using cultured isolates from the mice in this project by a nutrient dependency assay with L-cysteine indicating the use of these amino acids; however, validation of in vivo metabolism would indicate the biological host relevance. Understanding what drives these alternate metabolisms may prove invaluable for prevention and therapeutic development against chronic inflammatory disorders like ulcerative colitis and IBDs.

## Materials and Methods

### Sample collection and processing

#### IL-10 KO Mice

All mice in this study were IL10 KO Jax stock #002251/ C57BL/6J genetic background. Mice were bred in the Kansas State University Johnson Cancer Research Center mouse facility under *Helicobacter hepaticus-free* conditions. Figure 1 shows the experimental setup and treatment protocol of the study. To investigate the impact of the microbial populations on inherited dysbiosis, we conducted using four different experiment sets of mice - Control mice (Control); mice (with inherited dysbiosis) that were gavaged with control mice microbiota (FMT); mice (with inherited dysbiosis) that were gavaged with PBS (PBS); and mice (with inherited dysbiosis) that did not receive any gavage (No gavage). In order to instill inherited dysbiosis, we supplemented dams with autoclaved drinking water with cefoperazone sodium salt (CPZ; 0.5 mg/mL), a broad spectrum antibiotic, when they were in the third week of gestation. The duration of CPZ exposure lasted until pups were weaned at 21 days of age. We collected fecal samples from control mice daily, and were frozen to −80°C. To prepare the slurry for gavaging, we thawed, pooled, and gently vortexed control fecal samples in sterile PBS, then immediately frozen to −80°C until use. We gavaged the mice either with control mice slurry (FMT) or PBS (PBS) at 2, 3, and 8 weeks of age. Mice were monitored for signs of colitis. By week 23, remaining mice received 2% DSS for 5 days in autoclaved drinking water, and was changed as needed. We monitored mice daily during the time of DSS treatment and after treatment, until the end of the study. After a 16 day recovery period after 3% DSS treatment, all mice were sacrificed and tissues were collected.

#### Ethics Statement

All mouse experiments were reviewed and approved by the Institutional Animal Care and Use Committee at Kansas State University (APN #4391).

#### Serum

We collected blood via heart puncture, and centrifuged to collect serum. Serum was stored in −80°C until further analysis. Mouse Cytokine were obtained using a 31-Plex assay (Eve Technologies, Canada, catalog: #33836) that measured Eotaxin, G-CSF, GM-CSF, IFNγ, IL-1α, IL-1β, IL-2, IL-3, IL-4, IL-5, IL-6, IL-7, IL-9, IL-10, IL-12 (p40), IL-12 (p70), IL-13, IL-15, IL-17A, IP-10, KC, LIF, LIX, MCP-1, M-CSF, MIG, MIP-1α, MIP-1β, MIP-2, RANTES, TNFα, and VEGF-A (not validated).

#### Histopathology

A representative section of the lower colon from each mouse was routinely fixed in 10% neutral-buffered formalin, trimmed, embedded in paraffin, sectioned, and then stained with hematoxylin and eosin. Tissues were scored for characteristics of inflammation including surface epithelial damage, type and number of infiltrating inflammatory cells, and gland damage including dilation, inflammation, and hyperplasia. Each category was graded on a scale from 0 (normal), 1 (mild), 2 (moderate), or 3 (severe). Summative scores for each mouse and means for each treatment group were calculated.

#### Shotgun Metagenomics

We collected the fecal samples (n=51) and immediately froze to −80°C until extraction was performed. We extracted the gDNA using the E.Z.N.A Stool DNA Kit, following the manufacturer protocol for Pathogen detection supplemented with 10 minutes of bead beating. Extracted DNA was kept at −20°C until shotgun metagenomic sequencing. Sample DNA concentration and quality were determined by Qubit and Nanodrop. DNA libraries were constructed using Illumina’s Nextera XT Index Kit v2 kit. The products were then visualized on an Agilent Tapestation 4200 and size-selected using the BluePippin. The final library pool of 51 samples was quantified on the Kapa Biosystems qPCR protocol, and sequenced on the Illumina NovaSeq S1 chip in a 2 × 150bp paired-end sequencing run.

#### Colon RNA Sequencing

A small portion of the lower large intestine was stored in TRIzol reagent (−80°C) until sequencing. Samples were sent to CD Genomics (Shirley, New York), where they were extracted and QC analysis performed by manufacturer protocols. Next, cDNA library preparation (poly-A selection) was completed for the 12 samples, and were then sequenced via NovaSeq PE150 for a total of 20 million pair-end reads.

#### Bacterial Culturing and Nutrient Dependency Assay

Enteric bacteria from −80°C frozen mouse fecal samples from each treatment group were isolated on MacConkey Agar. Fecal pellets were vigorously homogenized in sterile PBS and streaking serial dilutions up to 10^-5^, incubating at 37°C overnight. Isolation streaks were performed twice for each isolate and DNA extracted using the E.Z.N.A Stool DNA Kit. Extracted DNA was sent out for Sanger sequencing by Functional Biosciences (Madison, Wisconsin). The bacterial growth curves were performed using M9 broth (Sigma) supplemented with magnesium chloride, calcium chloride, and 20% glucose. Media was supplemented with 100uM of cysteine and growth was monitored every 4 hours for 12 hours by OD600 measurements.

### Bioinformatic workflow

#### Metagenomics

We automated our metagenomics bioinformatics workflows using the program ‘anvi-run-workflow’ in anvi’o (70). The workflows use Snakemake (71), and implemented numerous tasks including short-read quality filtering, assembly, gene calling, functional annotation, hidden Markov model search (72), metagenomic read-recruitment and binning (73).

We used the program ‘iu-filer-quality-minoche’ to process the short metagenomic reads, and removed low-quality reads according to the criteria outlined by Minoche et al. We used MEGAHIT v1.2.9 (74) to co-assemble quality-filtered short reads into longer contiguous sequences (contigs). We assigned the metagenomes into 4 experimental groups (control/no-CPZ, n= 19; CPZ/no-gavage, n= 9; CPZ/gavage-with-PBS, n= 13; CPZ/FMT, n= 10) for the co-assembly. We then used the following strategies to process the assemblies: we used (1) ‘anvi-gen-contigs-database’ on contigs to compute k-mer frequencies and identify open reading frames (ORFs) using Prodigal v2.6.3 (75); (2) ‘anvi-run-hmms’ to identify sets of bacterial and archaeal single-copy core genes using HMMER v.3.2.1 (76); (3) ‘anvi-run-ncbi-cogs’ to annotate ORFs from NCBI’s Clusters of Orthologous Groups (COGs) (77); and (4) ‘anvi-run-kegg-kofams’ to annotate ORFs from KOfam HMM databases of KEGG orthologs (78). We mapped metagenomic short reads to contigs using Bowtie2 v2.3.5 (79) and converted to BAM files using samtools v1.9 (80). We profiled the BAM files using ‘anvi-profile’ with minimum contig length of 1000bp. We used ‘anvi-merge’ to combine all profiles into an anvi’o merged profile for all downstream analyses. We then used ‘anvi-cluster-contigs’ to group contigs into initial bins using CONCOCT v1.1.0 (81), and used ‘anvi-refine’ to manually curate the bins based on tetranucleotide frequency and different coverage across the samples. We marked bins that were more than 70% complete and less than 10% redundant as metagenome-assembled genomes (MAGs). Finally, we used ‘anvi-compute-genome-similarity’ to calculate average nucleotide identity (ANI) of our genomes using PyANI v0.2.9 (82), and identified non-redundant MAGs. We conducted all downstream analyses on only non-redundant MAGs. We used the “detection” metric to assess the occurrence of MAGs in a given sample. Detection of the MAGs is the proportion of the nucleotides that were covered by at least one short read. We considered a MAG was detected in a metagenome if the detection value was > 0.25, which is an appropriate cutoff to eliminate false-positive signals in read recruitment results, for its genome.

#### PATRIC analyses

Non-redundant genomes were uploaded and annotated in Pathosystems Resource Integration Center (PATRIC) (53). For the MAGs of interest, similar genomes were identified and phylogenetic trees were built using 50 similar strains to identify close relationships and approximate identity. Next, for each MAG, Subsystem genes were utilized for comparison of potential functions and virulence factors between the MAGs of interest. Counts of genes by each class of interest were used to build a heatmap and pathway constructs for each MAG of interest by using the Pathway function in PATRIC linked to KEGG.

#### RNASeq analysis

We used FastQC (83) and multiQC (84) to check the raw reads’ quality, and usedTrimmomatic (85) to trim the reads. We used Trinity ver 2.13.1 (86) for de novo assembly, and RSEM (87) and DESeq2 (88) to estimate the expression levels and differential expression analysis respectively. Annotation was carried out using blast+ 2.11 and R version 4.0.0. The transcripts compared in DESeq2 were first blasted against mouse transcript with C57BL/6NJ as a reference ensemble ID (http://ftp.ensembl.org/pub/release-107/fasta/mus_musculus_c57bl6nj/cdna/), then the top hit (lowest e-value and largest length with minimum of 150bp) were kept. Finally, we used GO_MWU (89) to analyze the annotated transcripts.

#### Statistical analyses

Differences between groups were analyzed by unpaired Student’s t-test, unless stated otherwise. P-values of less than 0.05 were considered statistically significant. Statistical analyses were performed in GraphPad Prism 9 or RStudio. We used the program - Data Integration Analysis for Biomarker discovery using Latent cOmponents (DIABLO) to build a model to identify key components in the metagenomic data and host RNASeq data (54, 90). We (i) first identified key omics variables in the integration process; (ii) and then maximized the common and correlated information between the datasets; and (iii) finally visualized the results to identify relevant microbial and host genomic markers.

### Data availability

#### Accession Number

Raw sequencing data for the mice metagenomes were uploaded to SRA under NCBI BioProject PRJNA843079. All other analyzed data in the form of databases and fasta files are accessible at figshare 10.6084/m9.figshare.21753623.

## Acknowledgements

We would like to acknowledge the services and help of Dr. Tracy Meisner for technique and experimental design. We appreciate Clark Bloomer and team with the Genome Sequencing Facility at the University of Kansas Medical Center for help with shotgun metagenomic sequencing. In addition, we would like to thank Dr. Sherry Fleming with the Johnson Cancer Center at Kansas State University for support and helpful discussion. We also acknowledge the Johnson Cancer Center at Kansas State University for funding and support for this project and personnel. Finally, we would like to acknowledge the 52 mice included in this study for their sacrifice to build our understanding of complex diseases such as IBD.

## Author Contributions

T.G.R. and S.T.M.L. designed the study. Sample collection was performed by T.G.R., L.H., A.K., S.P., K.R., T.S., H.W., and S.S. T.S., T.G.R., L.H., and A.K. completed DNA extraction with Nanodrop and Qubit quality analysis. Q.R. performed RNAseq bioinformatic analysis. T.G.R. and S.T.M.L. performed anvi’o and PATRIC bioinformatic analyses. Bacterial culturing and isolation was done by T.G.R. and S.S. T.G.R. and S.T.M.L. attributed biological relevance, wrote the manuscript, prepared figures, and supplementary files. T.G.R., S.T.M.L., and B.L.P. performed major manuscript and figure refinement while remaining authors contributed to lighter refinement. B.L.P. performed histologic analyses. S.T.M.L. acquired fundings for this study. All authors read, contributed to manuscript revision, and approved the submitted version.

## Supplemental Legends

Supplementary Table S1. Metagenomic sequencing results and pairings of all mouse samples.

Supplementary Table S2. Mouse histology scores, serum cytokine results and weights for each pup throughout the experiment.

Supplementary Table S3. Fold changes by treatment group comparison of select host genes.

Supplementary Table S4. Identities and taxonomies of all non-redundant MAGs.

Supplementary Table S5. Matrix of all non-redundant MAG detections in each pup.

Supplementary Figure S1. Rank-based Gene Ontology Analysis with Adaptive Clustering of control and FMT groups.

Supplementary Figure S2. Rank-based Gene Ontology Analysis with Adaptive Clustering of control and no gavage groups.

Supplementary Figure S3. Rank-based Gene Ontology Analysis with Adaptive Clustering of control and PBS groups.

Supplementary Figure S4. Rank-based Gene Ontology Analysis with Adaptive Clustering of FMT and no gavage groups.

Supplementary Figure S5. Rank-based Gene Ontology Analysis with Adaptive Clustering of FMT and PBS groups.

Supplementary Figure S6. Rank-based Gene Ontology Analysis with Adaptive Clustering of PBS and no gavage groups.

Supplementary Figure S7. Phylogenetic tree of the highest related strains of Enterobacteriaceae MAG 166.

Supplementary Figure S8. Phylogenetic tree of the highest related strains of Enterobacteriaceae MAG 169.

Supplementary Figure S9. Phylogenetic tree of the highest related strains of *Enterococcus* MAG 165.

Supplementary Figure S10. Phylogenetic tree of the highest related strains of *Akkermansia* MAG 001.

Supplementary Figure S11. Phylogenetic tree of the highest related strains of *Enterococcus* MAG 168.

Supplementary Figure S12. Phylogenetic tree of the highest related strains of Enterobacteriaceae MAG 167.

Supplementary Figure S13. Phylogenetic tree of the highest related strains of *Akkermansia* MAG 002.

## References

1. GBD 2017 Inflammatory Bowel Disease Collaborators. 2020. The global, regional, and national burden of inflammatory bowel disease in 195 countries and territories, 1990-2017:a systematic analysis for the Global Burden of Disease Study 2017. Lancet Gastroenterol Hepatol 5:17–30.

2. Ananthakrishnan AN, Bernstein CN, Iliopoulos D, Macpherson A, Neurath MF, Ali RAR, Vavricka SR, Fiocchi C. 2018. Environmental triggers in IBD: a review of progress and evidence. Nat Rev Gastroenterol Hepatol 15:39–49.

3. de Souza HSP, Fiocchi C. 2016. Immunopathogenesis of IBD: current state of the art. Nat Rev Gastroenterol Hepatol 13:13–27.

4. Chu H, Khosravi A, Kusumawardhani IP, Kwon AHK, Vasconcelos AC, Cunha LD, Mayer AE, Shen Y, Wu W-L, Kambal A, Targan SR, Xavier RJ, Ernst PB, Green DR, McGovern DPB, Virgin HW, Mazmanian SK. 2016. Gene-microbiota interactions contribute to the pathogenesis of inflammatory bowel disease. Science 352:1116–1120.

5. Wilks. Morbid appearances in the intestine of Miss Bankes. Med Times Gazette.

6. Mirsepasi-Lauridsen HC, Vallance BA, Krogfelt KA, Petersen AM. 2019. Escherichia coli Pathobionts Associated with Inflammatory Bowel Disease. Clin Microbiol Rev 32.

7. Devkota S, Wang Y, Musch MW, Leone V, Fehlner-Peach H, Nadimpalli A, Antonopoulos DA, Jabri B, Chang EB. 2012. Dietary-fat-induced taurocholic acid promotes pathobiont expansion and colitis in Il10-/-mice. Nature 487:104–108.

8. Yang H, Mirsepasi-Lauridsen HC, Struve C, Allaire JM, Sivignon A, Vogl W, Bosman ES, Ma C, Fotovati A, Reid GS, Li X, Petersen AM, Gouin SG, Barnich N, Jacobson K, Yu HB, Krogfelt KA, Vallance BA. 2020. Ulcerative Colitis-associated E. coli pathobionts potentiate colitis in susceptible hosts. Gut Microbes 12:1847976.

9. Bücker R, Schulz E, Günzel D, Bojarski C, Lee I-FM, John LJ, Wiegand S, Janßen T, Wieler LH, Dobrindt U, Beutin L, Ewers C, Fromm M, Siegmund B, Troeger H, Schulzke J-D. 2014. α-Haemolysin of Escherichia coli in IBD: a potentiator of inflammatory activity in the colon. Gut 63:1893–1901.

10. Renouf MJ, Cho YH, McPhee JB. 2019. Emergent Behavior of IBD-Associated Escherichiacoli During Disease. Inflamm Bowel Dis 25:33–44.

11. Golińska E, Tomusiak A, Gosiewski T, Więcek G, Machul A, Mikołajczyk D, Bulanda M, Heczko PB, Strus M. 2013. Virulence factors of Enterococcus strains isolated from patients with inflammatory bowel disease. World J Gastroenterol 19:3562–3572.

12. Cameron EA, Sperandio V, Dunny GM. 2019. Enterococcus faecalis Enhances Expression and Activity of the Enterohemorrhagic Escherichia coli Type III Secretion System. MBio 10.

13. Kang C-S, Ban M, Choi E-J, Moon H-G, Jeon J-S, Kim D-K, Park S-K, Jeon SG, Roh T-Y, Myung S-J, Gho YS, Kim JG, Kim Y-K. 2013. Extracellular vesicles derived from gut microbiota, especially Akkermansia muciniphila, protect the progression of dextran sulfate sodium-induced colitis. PLoS One 8:e76520.

14. Reunanen J, Kainulainen V, Huuskonen L, Ottman N, Belzer C, Huhtinen H, de Vos WM, Satokari R. 2015. Akkermansia muciniphila Adheres to Enterocytes and Strengthens the Integrity of the Epithelial Cell Layer. Appl Environ Microbiol 81:3655–3662.

15. Geirnaert A, Calatayud M, Grootaert C, Laukens D, Devriese S, Smagghe G, De Vos M, Boon N, Van de Wiele T. 12/2017. Butyrate-producing bacteria supplemented in vitro to Crohn’s disease patient microbiota increased butyrate production and enhanced intestinal epithelial barrier integrity. Sci Rep 7:11450.

16. Baldelli V, Scaldaferri F, Putignani L, Del Chierico F. 2021. The Role of Enterobacteriaceae in Gut Microbiota Dysbiosis in Inflammatory Bowel Diseases. Microorganisms 9.

17. Spees AM, Wangdi T, Lopez CA, Kingsbury DD, Xavier MN, Winter SE, Tsolis RM, Bäumler AJ. 2013. Streptomycin-induced inflammation enhances Escherichia coli gut colonization through nitrate respiration. MBio 4.

18. Winter SE, Winter MG, Xavier MN, Thiennimitr P, Poon V, Keestra AM, Laughlin RC, Gomez G, Wu J, Lawhon SD, Popova IE, Parikh SJ, Adams LG, Tsolis RM, Stewart VJ, Baumler AJ. 2013. Host-Derived Nitrate Boosts Growth of E. coli in the Inflamed Gut. Science 339:708–711.

19. Wang J, Guo X, Li H, Qi H, Qian J, Yan S, Shi J, Niu W. 2019. Hydrogen Sulfide From Cysteine Desulfurase, Not 3-Mercaptopyruvate Sulfurtransferase, Contributes to Sustaining Cell Growth and Bioenergetics in E. coli Under Anaerobic Conditions. Front Microbiol 10:2357.

20. Walker A, Schmitt-Kopplin P. 2021. The role of fecal sulfur metabolome in inflammatory bowel diseases. Int J Med Microbiol 311:151513.

21. Rezaie A, Parker RD, Abdollahi M. 2007. Oxidative stress and pathogenesis of inflammatory bowel disease: an epiphenomenon or the cause? Dig Dis Sci 52:2015–2021.

22. Alemany-Cosme, Sáez-González, Moret. 2021. Oxidative stress in the pathogenesis of Crohn’s disease and the interconnection with immunological response, microbiota, external environmental factors, and … Antioxidants (Basel).

23. Henrick BM, Rodriguez L, Lakshmikanth T, Pou C, Henckel E, Arzoomand A, Olin A, Wang J, Mikes J, Tan Z, Chen Y, Ehrlich AM, Bernhardsson AK, Mugabo CH, Ambrosiani Y, Gustafsson A, Chew S, Brown HK, Prambs J, Bohlin K, Mitchell RD, Underwood MA, Smilowitz JT, German JB, Frese SA, Brodin P. 2021. Bifidobacteria-mediated immune system imprinting early in life. Cell 184:3884–3898.e11.

24. Milani C, Duranti S, Bottacini F, Casey E, Turroni F, Mahony J, Belzer C, Delgado Palacio S, Arboleya Montes S, Mancabelli L, Lugli GA, Rodriguez JM, Bode L, de Vos W, Gueimonde M, Margolles A, van Sinderen D, Ventura M. 2017. The First Microbial Colonizers of the Human Gut: Composition, Activities, and Health Implications of the Infant Gut Microbiota. Microbiol Mol Biol Rev 81.

25. Torow N, Hornef MW. 2017. The Neonatal Window of Opportunity: Setting the Stage for Life-Long Host-Microbial Interaction and Immune Homeostasis. J Immunol 198:557–563.

26. Soufli I, Toumi R, Rafa H, Touil-Boukoffa C. 2016. Overview of cytokines and nitric oxide involvement in immuno-pathogenesis of inflammatory bowel diseases. World J Gastrointest Pharmacol Ther 7:353–360.

27. Torrence AE, Brabb T, Viney JL, Bielefeldt-Ohmann H, Treuting P, Seamons A, Drivdahl R, Zeng W, Maggio-Price L. 2008. Serum biomarkers in a mouse model of bacterial-induced inflammatory bowel disease. Inflamm Bowel Dis 14:480–490.

28. Rocha BS, Laranjinha J. 2020. Nitrate from diet might fuel gut microbiota metabolism:Minding the gap between redox signaling and inter-kingdom communication. Free Radic Biol Med 149:37–43.

29. Schmitt H, Neurath MF, Atreya R. 2021. Role of the IL23/IL17 Pathway in Crohn’s Disease. Front Immunol 12:622934.

30. Kushkevych I, Dordević D, Vítězová M. 2021. Possible synergy effect of hydrogen sulfide and acetate produced by sulfate-reducing bacteria on inflammatory bowel disease development. J Advert Res 27:71–78.

31. Porrini C, Ramarao N, Tran S-L. 2020. Dr. NO and Mr. Toxic - the versatile role of nitric oxide. Biol Chem 401:547–572.

32. Baranipour S, Amini Kadijani A, Qujeq D, Shahrokh S, Haghazali M, Mirzaei A, Asadzadeh-Aghdaei H. 2018. Inducible nitric oxide synthase as a potential blood-based biomarker in inflammatory bowel diseases. Gastroenterol Hepatol Bed Bench 11:S124–S128.

33. Tarling EJ, de Aguiar Vallim TQ, Edwards PA. 2013. Role of ABC transporters in lipid transport and human disease. Trends Endocrinol Metab 24:342–350.

34. Ferrer-Picón E, Dotti I, Corraliza AM, Mayorgas A, Esteller M, Perales JC, Ricart E, Masamunt MC, Carrasco A, Tristán E, Esteve M, Salas A. 2020. Intestinal Inflammation Modulates the Epithelial Response to Butyrate in Patients With Inflammatory Bowel Disease. Inflamm Bowel Dis 26:43–55.

35. Fransén K, Klintenäs M, Osterström A, Dimberg J, Monstein H-J, Söderkvist P. 2004. Mutation analysis of the BRAF, ARAF and RAF-1 genes in human colorectal adenocarcinomas. Carcinogenesis 25:527–533.

36. Watson SF, Bellora N, Macias S. 2020. ILF3 contributes to the establishment of the antiviral type I interferon program. Nucleic Acids Res 48:116–129.

37. Bubier JA, Philip VM, Quince C, Campbell J, Zhou Y. 2018. Systems genetic discovery of host-microbiome interactions reveals mechanisms of microbial involvement in disease. bioRxiv.

38. van der Graaf A, Zorro MM, Claringbould A, Võsa U, Aguirre-Gamboa R, Li C, Mooiweer J, Ricaño-Ponce I, Borek Z, Koning F, Kooy-Winkelaar Y, Sollid LM, Qiao S-W, Kumar V, Li Y, Franke L, Withoff S, Wijmenga C, Sanna S, Jonkers I, Bios Consortium. 2021. Systematic Prioritization of Candidate Genes in Disease Loci Identifies TRAFD1 as a Master Regulator of IFNγSignaling in Celiac Disease. Front Genet 11.

39. Sido B, Hack V, Hochlehnert A, Lipps H, Herfarth C, Dröge W. 1998. Impairment of intestinal glutathione synthesis in patients with inflammatory bowel disease. Gut 42:485–492.

40. Morgenstern I, Raijmakers MTM, Peters WHM, Hoensch H, Kirch W. 2003. Homocysteine,cysteine, and glutathione in human colonic mucosa: elevated levels of homocysteine in patients with inflammatory bowel disease. Dig Dis Sci 48:2083–2090.

41. Membrillo-Hernández J, Coopamah MD, Channa A, Hughes MN, Poole RK. 1998. A novel mechanism for upregulation of the Escherichia coli K-12 hmp (flavohaemoglobin) gene by the “NO releaser”, S-nitrosoglutathione: nitrosation of homocysteine and modulation of MetR binding to the glyA-hmp intergenic region. Mol Microbiol 29:1101–1112.

42. Chassaing B, Koren O, Carvalho FA, Ley RE, Gewirtz AT. 2014. AIEC pathobiont instigates chronic colitis in susceptible hosts by altering microbiota composition. Gut 63:1069–1080.

43. Boland K, Bedrani L, Turpin W, Kabakchiev B, Stempak J, Borowski K, Nguyen G, Steinhart AH, Smith MI, Croitoru K, Silverberg MS. 2021. Persistent Diarrhea in Patients With Crohn’s Disease After Mucosal Healing Is Associated With Lower Diversity of the Intestinal Microbiome and Increased Dysbiosis. Clin Gastroenterol Hepatol 19:296–304.e3.

44. Campbell JH, O’Donoghue P, Campbell AG, Schwientek P, Sczyrba A, Woyke T, Söll D, Podar M. 2013. UGA is an additional glycine codon in uncultured SR1 bacteria from the human microbiota. Proc Natl Acad Sci U S A 110:5540–5545.

45. Bowers RM, Kyrpides NC, Stepanauskas R, Harmon-Smith M, Doud D, Reddy TBK, Schulz F, Jarett J, Rivers AR, Eloe-Fadrosh EA, Tringe SG, Ivanova NN, Copeland A, Clum A, Becraft ED, Malmstrom RR, Birren B, Podar M, Bork P, Weinstock GM, Garrity GM, Dodsworth JA, Yooseph S, Sutton G, Glöckner FO, Gilbert JA, Nelson WC, Hallam SJ, Jungbluth SP, Ettema TJG, Tighe S, Konstantinidis KT, Liu W-T, Baker BJ, Rattei T, Eisen JA, Hedlund B, McMahon KD, Fierer N, Knight R, Finn R, Cochrane G, Karsch-Mizrachi I, Tyson GW, Rinke C, Genome Standards Consortium, Lapidus A, Meyer F, Yilmaz P, Parks DH, Eren AM, Schriml L, Banfield JF, Hugenholtz P, Woyke T. 2017. Minimum information about a single amplified genome (MISAG) and a metagenome-assembled genome(MIMAG) of bacteria and archaea. Nat Biotechnol 35:725–731.

46. Sharon I, Morowitz MJ, Thomas BC, Costello EK, Relman DA, Banfield JF. 2013. Time series community genomics analysis reveals rapid shifts in bacterial species, strains, and phage during infant gut colonization. Genome Res 23:111–120.

47. Lee STM, Kahn SA, Delmont TO, Shaiber A, Esen ÖC, Hubert NA, Morrison HG, Antonopoulos DA, Rubin DT, Eren AM. 2017. Tracking microbial colonization in fecal microbiota transplantation experiments via genome-resolved metagenomics. Microbiome 5:50.

48. Garrett WS, Gallini CA, Yatsunenko T, Michaud M, DuBois A, Delaney ML, Punit S, Karlsson M, Bry L, Glickman JN, Gordon JI, Onderdonk AB, Glimcher LH. 2010. Enterobacteriaceae act in concert with the gut microbiota to induce spontaneous and maternally transmitted colitis. Cell Host Microbe 8:292–300.

49. Zhou Y, Chen H, He H, Du Y, Hu J, Li Y, Li Y, Zhou Y, Wang H, Chen Y, Nie Y. 2016. Increased Enterococcus faecalis infection is associated with clinically active Crohn disease. Medicine 95:e5019.

50. Pham HN, Ohkusu K, Mishima N, Noda M, Monir Shah M, Sun X, Hayashi M, Ezaki T. 2007. Phylogeny and species identification of the family Enterobacteriaceae based on dnaJ sequences. Diagn Microbiol Infect Dis 58:153–161.

51. Janda JM, Abbott SL. 2007. 16S rRNA gene sequencing for bacterial identification in the diagnostic laboratory: pluses, perils, and pitfalls. J Clin Microbiol 45:2761–2764.

52. Ondov BD, Treangen TJ, Melsted P, Mallonee AB, Bergman NH, Koren S, Phillippy AM. 2016. Mash: fast genome and metagenome distance estimation using MinHash. Genome Biol 17:132.

53. Wattam AR, Davis JJ, Assaf R, Boisvert S, Brettin T, Bun C, Conrad N, Dietrich EM, Disz T, Gabbard JL, Gerdes S, Henry CS, Kenyon RW, Machi D, Mao C, Nordberg EK, Olsen GJ, Murphy-Olson DE, Olson R, Overbeek R, Parrello B, Pusch GD, Shukla M, Vonstein V, Warren A, Xia F, Yoo H, Stevens RL. 2017. Improvements to PATRIC, the all-bacterial Bioinformatics Database and Analysis Resource Center. Nucleic Acids Res 45:D535–D542.

54. Rohart F, Gautier B, Singh A, Lê Cao K-A. 2017. mixOmics: An R package for ‘omics feature selection and multiple data integration. PLoS Comput Biol 13:e1005752.

55. Park E, Park SY, Wang C, Xu J, LaFauci G, Schuller-Levis G. 2002. Cloning of murine cysteine sulfinic acid decarboxylase and its mRNA expression in murine tissues11The sequence presented in this paper is available from GenBank with the accession number AY033912. Biochimica et Biophysica Acta (BBA) - Gene Structure and Expression 1574:403–406.

56. Kim CJ, Kovacs-Nolan J, Yang C, Archbold T, Fan MZ, Mine Y. 2009. L-cysteine supplementation attenuates local inflammation and restores gut homeostasis in a porcine model of colitis. Biochim Biophys Acta 1790:1161–1169.

57. Xiao JY. 2015. Nutritional interventions during recovery in protein deficient piglets with dextran sulfate sodium induced colitis. search.proquest.com.

58. Bian X, Wu W, Yang L, Lv L, Wang Q, Li Y, Ye J, Fang D, Wu J, Jiang X, Shi D, Li L. 2019. Administration of Akkermansia muciniphila Ameliorates Dextran Sulfate Sodium-Induced Ulcerative Colitis in Mice. Front Microbiol 10:2259.

59. Ottman N, Reunanen J, Meijerink M, Pietilä TE, Kainulainen V, Klievink J, Huuskonen L, Aalvink S, Skurnik M, Boeren S, Satokari R, Mercenier A, Palva A, Smidt H, de Vos WM, Belzer C. 2017. Pili-like proteins of Akkermansia muciniphila modulate host immune responses and gut barrier function. PLoS One 12:e0173004.

60. Wang S, El-Fahmawi A, Christian DA, Fang Q, Radaelli E, Chen L, Sullivan MC, Misic AM, Ellringer JA, Zhu X-Q, Winter SE, Hunter CA, Beiting DP. 2019. Infection-Induced Intestinal Dysbiosis Is Mediated by Macrophage Activation and Nitrate Production. MBio 10:e00935–19, /mbio/10/3/mBio.00935-19.atom.

61. Stipanuk MH. 2020. Metabolism of Sulfur-Containing Amino Acids: How the Body Copes with Excess Methionine, Cysteine, and Sulfide. J Nutr 150:2494S–2505S.

62. Carbonero F, Benefiel AC, Alizadeh-Ghamsari AH, Gaskins HR. 2012. Microbial pathways in colonic sulfur metabolism and links with health and disease. Front Physiol 3:448.

63. Kumar P, Liu C, Hsu JW, Chacko S, Minard C, Jahoor F, Sekhar RV. 2021. Glycine and N-acetylcysteine (GlyNAC) supplementation in older adults improves glutathione deficiency,oxidative tress, mitochondrial dysfunction, inflammation, insulin resistance, endothelial dysfunction, genotoxicity, muscle strength, and cognition: Results of a pilot clinical trial. Clin Transl Med 11:e372.

64. Singh H, Dai Y, Outten FW, Busenlehner LS. 2013. Escherichia coli SufE sulfur transfer protein modulates the SufS cysteine desulfurase through allosteric conformational dynamics. J Biol Chem 288:36189–36200.

65. Dunkle JA, Bruno MR, Frantom PA. 2020. Structural evidence for a latch mechanism regulating access to the active site of SufS-family cysteine desulfurases. Acta Crystallogr D Struct Biol 76:291–301.

66. Nishikawa M, Shen L, Ogawa K’ ichi. 2018. Taurine dioxygenase (tauD)-independent taurine assimilation in Escherichia coli. Microbiology 164:1446–1456.

67. Riboldi GP, Larson TJ, Frazzon J. 2011. Enterococcus faecalis sufCDSUB complements Escherichia coli sufABCDSE. FEMS Microbiol Lett 320:15–24.

68. Reinders CA, Jonkers D, Janson EA, Stockbrügger RW, Stobberingh EE, Hellström PM, Lundberg JO. 2007. Rectal nitric oxide and fecal calprotectin in inflammatory bowel disease. Scand J Gastroenterol 42:1151–1157.

69. Tiso M, Schechter AN. 2015. Nitrate reduction to nitrite, nitric oxide and ammonia by gut bacteria under physiological conditions. PLoS One 10:e0119712.

70. Eren AM, Kiefl E, Shaiber A, Veseli I, Miller SE, Schechter MS, Fink I, Pan JN, Yousef M, Fogarty EC, Trigodet F, Watson AR, Esen ÖC, Moore RM, Clayssen Q, Lee MD, Kivenson V, Graham ED, Merrill BD, Karkman A, Blankenberg D, Eppley JM, Sjödin A, Scott JJ, Vázquez-Campos X, McKay LJ, McDaniel EA, Stevens SLR, Anderson RE, Fuessel J, Fernandez-Guerra A, Maignien L, Delmont TO, Willis AD. 2021. Community-led, integrated,reproducible multi-omics with anvi’o. Nat Microbiol 6:3–6.

71. Shaiber A, Willis AD, Delmont TO, Roux S, Chen L-X, Schmid AC, Yousef M, Watson AR, Lolans K, Esen ÖC, Lee STM, Downey N, Morrison HG, Dewhirst FE, Mark Welch JL, Eren AM. 2020. Functional and genetic markers of niche partitioning among enigmatic members of the human oral microbiome. Genome Biol 21:292.

72. Johnson LS, Eddy SR, Portugaly E. 2010. Hidden Markov model speed heuristic and iterative HMM search procedure. BMC Bioinformatics 11:431.

73. Murat Eren A, Esen ÖC, Quince C, Vineis JH, Morrison HG, Sogin ML, Delmont TO. 2015. Anvi’o: an advanced analysis and visualization platform for ‘omics data. PeerJ 3:e1319.

74. Li D, Liu C-M, Luo R, Sadakane K, Lam T-W. 2015. MEGAHIT: an ultra-fast single-node solution for large and complex metagenomics assembly via succinct de Bruijn graph. Bioinformatics 31:1674–1676.

75. Hyatt D, Chen G-L, Locascio PF, Land ML, Larimer FW, Hauser LJ. 2010. Prodigal:prokaryotic gene recognition and translation initiation site identification. BMC Bioinformatics 11:119.

76. Wheeler TJ, Eddy SR. 2013. nhmmer: DNA homology search with profile HMMs. Bioinformatics 29:2487–2489.

77. Galperin MY, Wolf YI, Makarova KS, Vera Alvarez R, Landsman D, Koonin EV. 2021. COG database update: focus on microbial diversity, model organisms, and widespread pathogens. Nucleic Acids Res 49:D274–D281.

78. Aramaki T, Blanc-Mathieu R, Endo H, Ohkubo K, Kanehisa M, Goto S, Ogata H. 2020. KofamKOALA: KEGG Ortholog assignment based on profile HMM and adaptive score threshold. Bioinformatics 36:2251–2252.

79. Langmead B, Salzberg SL. 2012. Fast gapped-read alignment with Bowtie 2. Nat Methods 9:357–359.

80. Danecek P, Bonfield JK, Liddle J, Marshall J, Ohan V, Pollard MO, Whitwham A, Keane T, McCarthy SA, Davies RM, Li H. 2021. Twelve years of SAMtools and BCFtools. Gigascience 10.

81. Alneberg J, Bjarnason BS, de Bruijn I, Schirmer M, Quick J, Ijaz UZ, Loman NJ, Andersson AF, Quince C. 2013. CONCOCT: Clustering cONtigs on COverage and ComposiTion. arXiv [q-bioGN].

82. Pritchard L, Glover RH, Humphris S, Elphinstone JG, Toth IK. 2015. Genomics and taxonomy in diagnostics for food security: soft-rotting enterobacterial plant pathogens. Anal Methods 8:12–24.

83. Andrews S. FastQC: A quality control analysis tool for high throughput sequencing data. Github. https://github.com/s-andrews/FastQC. Retrieved 22 September 2022.

84. Ewels P, Magnusson M, Lundin S, Käller M. 2016. MultiQC: summarize analysis results for multiple tools and samples in a single report. Bioinformatics 32:3047–3048.

85. Bolger AM, Lohse M, Usadel B. 2014. Trimmomatic: a flexible trimmer for Illumina sequence data. Bioinformatics 30:2114–2120.

86. Grabherr MG, Haas BJ, Yassour M, Levin JZ, Thompson DA, Amit I, Adiconis X, Fan L, Raychowdhury R, Zeng Q, Chen Z, Mauceli E, Hacohen N, Gnirke A, Rhind N, di Palma F, Birren BW, Nusbaum C, Lindblad-Toh K, Friedman N, Regev A. 2011. Full-length transcriptome assembly from RNA-Seq data without a reference genome. Nat Biotechnol 29:644–652.

87. Li B, Dewey CN. 2011. RSEM: accurate transcript quantification from RNA-Seq data with or without a reference genome. BMC Bioinformatics 12:323.

88. Love MI, Huber W, Anders S. 2014. Moderated estimation of fold change and dispersion for RNA-seq data with DESeq2. Genome Biol 15:550.

89. Wright RM, Aglyamova GV, Meyer E, Matz MV. 2015. Gene expression associated with white syndromes in a reef building coral, Acropora hyacinthus. BMC Genomics 16:371.

90. Kim-Anh Le Cao, Florian Rohart, Ignacio Gonzalez, Sebastien Dejean with key contributors Benoit Gautier, Francois Bartolo, contributions from Pierre Monget, Jeff Coquery, FangZou Yao and Benoit Liquet. 2016. mixOmics: Omics Data Integration Project. R package version 6.1.1.

